# Nitrate leaching and its implication for Fe and As mobility in a Southeast Asian aquifer

**DOI:** 10.1101/2022.10.07.511258

**Authors:** Martyna Glodowska, Yinxiao Ma, Garrett Smith, Andreas Kappler, Mike Jetten, Cornelia U. Welte

## Abstract

The drinking water quality of millions of people in South and Southeast Asia is at risk due to arsenic (As) contamination of groundwater and insufficient access to water treatment facilities. Intensive use of nitrogen (N) fertilizer increases the possibility of nitrate (NO_3_^-^) leaching into aquifers, yet very little is known about how the N cycle will interact with and affect the iron (Fe) and As mobility in aquifers. We hypothesized that input of NO_3_^-^ into highly methanogenic aquifers can stimulate nitrate-dependent anaerobic methane oxidation (N-DAMO) and subsequently help to remove NO_3_^-^ and decrease CH_4_ emission. We, therefore, investigated the effects of N input into aquifers and its effect on Fe and As mobility, by running a set of microcosm experiments using aquifer sediment from Van Phuc, Vietnam supplemented with ^15^NO_3_^-^ and ^13^CH_4_. Additionally, we assessed the effect of N-DAMO by inoculating the sediment with two different N-DAMO enrichment cultures (N-DAMO(O) and N-DAMO(V)). We found that native microbial communities and both N-DAMO enrichments could efficiently consume nearly 5 mM NO_3_^-^ in 5 days. In an uninoculated setup, NO_3_^-^ was preferentially used over Fe(III) as electron acceptor and consequently inhibited Fe(III) reduction and As mobilization. The addition of N-DAMO(O) and N-DAMO(V) enrichment cultures led to substantial Fe(III) reduction followed by the release of Fe^2+^ (0.190±0.002 mM and 0.350±0.007 mM, respectively) and buildup of sedimentary Fe(II) (11.20±0.20 mM and 10.91±0.47 mM, respectively) at the end of the experiment (day 64). Only in the N-DAMO(O) inoculated setup, As was mobilized (27.1±10.8 μg/L), while in the setup inoculated with N-DAMO(V) a significant amount of Mn (24.15±0.41 mg/L) was released to the water. Methane oxidation and ^13^CO_2_ formation were observed only in the inoculated setups, suggesting that the native microbial community did not have sufficient potential for N-DAMO. An increase of NH_4_^+^ implied that dissimilatory nitrate reduction to ammonium (DNRA) took place in both inoculated setups. The archaeal community in all treatments was dominated by *Ca*. Methanoperedens while the bacterial community consisted largely of various denitrifiers. Overall, our results suggest that input of N fertilizers to the aquifer decreases As mobility and that CH_4_ cannot serve as an electron donor for the native NO_3_^-^ reducing community.

**Graphical abstract:** 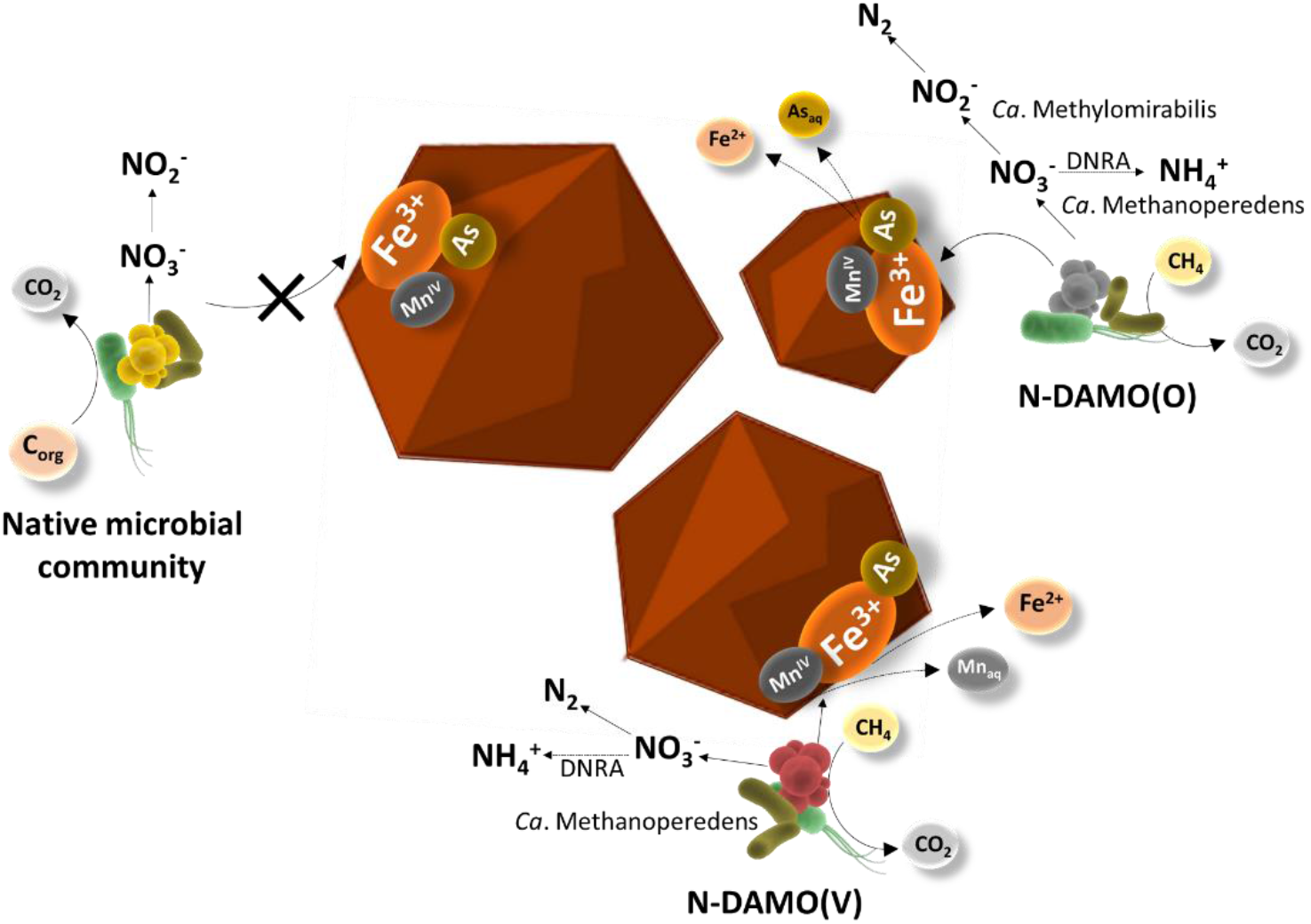

## Introduction

High arsenic (As) concentrations in groundwater are a global problem. It was estimated that as much as 150 million people worldwide might be affected by As-contaminated water exceeding the drinking water limit of 10 μg/L recommended by the World Health Organization (WHO), with the vast majority of affected people (~ 94%) located in Asia (Podgorski and Berg, 2020). Arsenic has been recognized as a group I human carcinogen by The International Agency for Research on Cancer (IARC). Long-term exposure to As-contaminated water or excessive As intake is a serious health hazard frequently leading to an increased risk of cancer, and cardiovascular and neurological diseases (Chen et al., 2009; Hughes, 2002). Therefore, As pollution has become an alarming concern triggering a global research initiative aiming to understand the underlying biogeochemical mechanisms of As (im)mobilization in aquifers. Due to limited access to water treatment facilities and use of the untreated shallow groundwater as a primary drinking water source, As poisoning is particularly severe in rural areas of South and Southeast Asia (Carrard et al., 2019). Vietnam is among the most affected countries where As concentrations in drinking water from household water wells can reach 3050 μg/L exceeding 300 times the WHO safe limit (Berg et al., 2007; Le Luu, 2019). The problem of high As concentration in drinking water has not been solved yet, and it continues to be the largest mass poisoning of the human population in history (Sen and Biswas, 2013).

Mobility of As in an aquifer is the main factor leading to its accumulation in water which is controlled by sediment geochemistry, evapotranspiration, flow-through conditions, pH, redox potential, microbial community, and ion availability (Mladenov et al., 2014; Pipattanajaroenkul et al., 2021). Because of the strong affinity of As and adsorption ability of Fe(III) (oxyhydr)oxides, the reductive dissolution of Fe(III) minerals plays an important role in As groundwater accumulation (Yang et al., 2015). The coupling of reductive dissolution of Fe(III) (oxyhydr)oxides with organic carbon oxidation by microbial processes is considered the primary pathway for increasing dissolved As concentrations in aquifers of South and Southeast Asia (Fendorf et al., 2010). Several studies showed that the presence of Fe(III)-reducing microorganisms significantly increased the rate of Fe(III)-reduction and As mobilization (Glodowska et al., 2020; Islam et al., 2005a, 2005b; Jiang et al., 2013). Arsenic is usually bound to the surface of Fe(III) (oxyhydr)oxide minerals in the form of As(V). When the Fe(III) mineral is reduced to dissolved or solid-phase Fe(II), As is also released from the Fe(III) minerals (Qiao et al., 2021). More crystalline Fe(III) minerals such as magnetite, goethite, or hematite, are generally less bioavailable for the microorganisms, and therefore are also less likely to release As. In contrast, ferrihydrite is a poorly crystalline mineral and thus more prone to reduction and more easily releases As into the water than other Fe(III) (oxyhydr)oxides (Das et al., 2014).

The nitrogen (N) cycle may change As mobility in groundwater by affecting the conversion of Fe(III) to Fe(II) (Fig. 1). Nitrogen is widely present in various environments, and its primary forms in groundwater are nitrate (NO_3_^-^) and ammonium (NH_4_^+^), while nitrite (NO_2_^-^) is found at relatively low concentrations or is absent (Parvizishad et al., 2017). Due to increasing agricultural production in past decades, the excessive use of fertilizer has introduced additional N (mainly NO_3_^-^ and NH_4_^+^) into soil and surface water, causing eutrophication in these environments (Conley et al., 2009). Furthermore, it has been shown that N fertilizers can leach into the groundwater in the form of NH_4_^+^ or NO_3_^-^, causing the total N content in the groundwater to increase (Bijay-Singh and Craswell, 2021). In the presence of oxygen (O_2_), NH_4_^+^ can be oxidized to NO_3_^-^ via nitrification (equation 1) by ammonia-oxidizing bacteria or archaea (Jetten et al., 1998; Könneke et al., 2005). Nitrification of NH_4_^+^ is the primary source of NO_3_^-^ in aquifers (Umezawa et al., 2008). In nitrification-dominated environments, when both NO_3_^-^ and Fe(III) are present in groundwater, heterotrophic microorganisms will likely preferentially utilize NO_3_^-^ as an electron acceptor due to the higher Gibbs free energy change (equations 2 and 3) (Hanson et al., 2013; Lovley and Phillips, 1988). Thus, the presence of NO_3_^-^ can inhibit the reduction of Fe(III) (oxyhydr)oxides, preventing As mobilization to the aquifer, which in its pentavalent form remains stably adsorbed to the Fe(III) mineral (Weng et al., 2017). Moreover, a previous study showed that the addition of NO_3_^-^ stimulates anoxic nitrate-dependent Fe(II) oxidation leading to a decrease in dissolved Fe(II) and As in groundwater (Harvey et al., 2002; Smith et al., 2017). This is because NO_3_^-^ can oxidize Fe^2+^ to Fe^3+^ via biotic or abiotic reactions simultaneously co-precipitating with dissolved As.

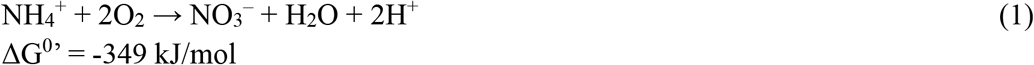

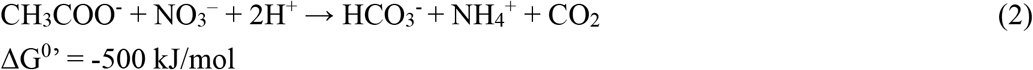

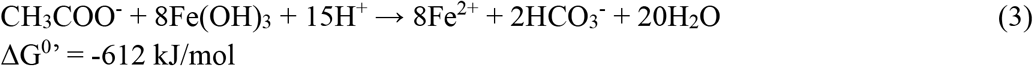

Due to the presence of clay minerals and overall net negative charge of soil and sediment particles, NO_3_^-^ is more easily transported by water flow into the subsurface aquifer compared to the positively charged NH_4_^+^ (Köhler et al., 2006; Nieder et al., 2011). Consequently, heterotrophic denitrification may take place in anoxic underground aquifers (equation 4) (Austin et al., 2016).

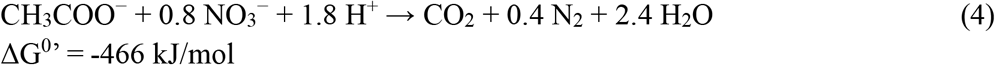

Denitrifying bacteria subsequently reduce NO_3_^-^ to NO_2_^-^, nitric oxide (NO), nitrous oxide (N_2_O), and ultimately to dinitrogen gas (N_2_). Denitrifiers can be either autotrophic or heterotrophic depending on the source of electrons. Heterotrophic denitrifying bacteria and archaea usually couple the oxidation of organic matter with NO_3_^-^ reduction. Autotrophic denitrifying bacteria however use NO_3_^-^ to oxidize inorganic reduced substances such as Fe(II) and Mn(II) (Kappler et al., 2021; Weber et al., 2006). During this process, Fe(II) is oxidized to Fe(III) to form poorly soluble Fe(III) (oxyhydr)oxides, which can later co-precipitate with As (Hohmann et al., 2010).

**Fig. 1.**
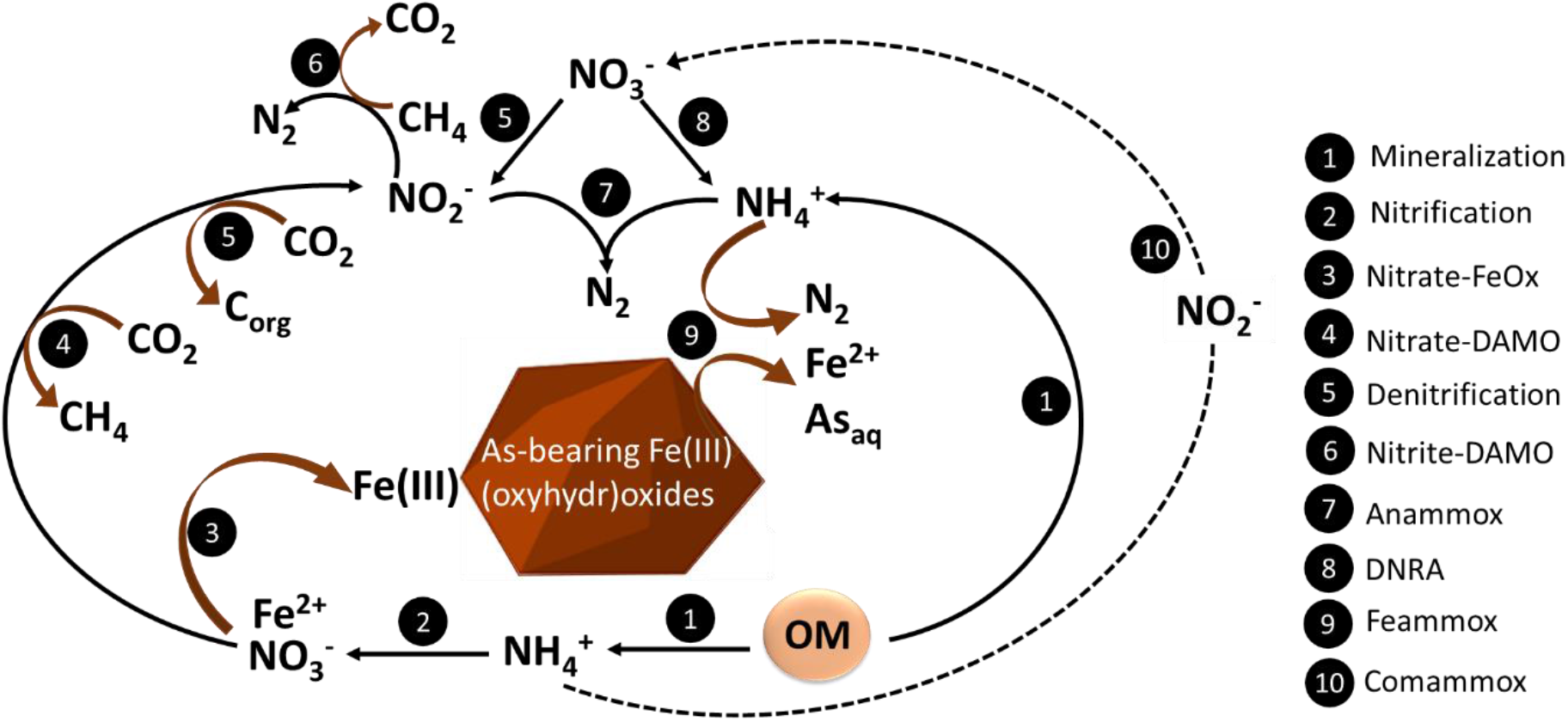
Possible nitrogen cycle reactions in an aquifer and their effect on Fe and As mobility. Nitrate-dependent Fe^2+^ oxidation (Nitrate-FeOx), dissimilatory nitrate reduction to ammonium (DNRA), nitrate/nitrite-dependent anaerobic methane oxidation (Nitrate/Nitrite-DAMO), nitrate-dependent Fe^2+^ oxidation (Nitrate-FeOx)

It was also shown that some microorganisms are capable of nitrate-dependent anaerobic methane oxidation (N-DAMO), a process mediated by ANME-2d archaea, specifically by *Candidatus* Methanoperedens (Haroon et al., 2013; Raghoebarsing et al., 2006). Since their discovery, several N-DAMO archaea have been enriched from anoxic freshwater sediments, digester sludge, and rice paddies (Arshad et al., 2015; Hu et al., 2009; Vaksmaa et al., 2017). The N-DAMO archaea reduce NO_3_^-^ to NO_2_^-^ while oxidizing methane (CH_4_) to gain energy (equation 5). Nitrite-dependent methanotrophic bacteria named *Candidatus* Methylomirabilis oxidize NO_2_^-^ to N_2_ at the expense of CH_4_ (equation 6) (Ettwig et al., 2010).

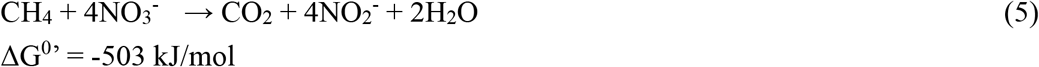

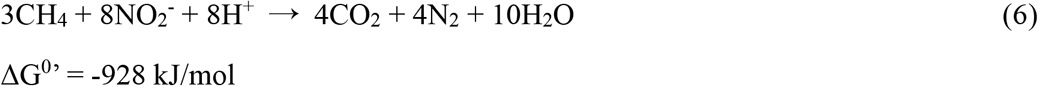

Together, the N-DAMO process might be particularly relevant for strongly methanogenic aquifers in agricultural areas where the intensive application of fertilizer leads to the NO_3_^-^ accumulation, however, to date N-DAMO activity has not been confirmed in aquifer systems.

Feammox is one of the newly proposed pathways coupling NH_4_^+^ oxidation with Fe(III) reduction (equation 7) that could potentially lead to As release. Previous studies have shown that indeed Fe(III)-reducing *Acidimicrobiaceae* -like bacteria can use NH_4_^+^ as an electron donor at low pH (Huang and Jaffé, 2018). Some studies have suggested that the Feammox process plays an important role in the N cycle in various ecosystems such as tropical forest soils, paddy fields, rivers, and lake sediments (Li et al., 2019; Rios-Del Toro et al., 2018). Although there have been some reports of the positive correlation between high NH_4_^+^ and high As in Southeast Asian aquifers until now it remains unclear whether NH_4_^+^ is involved in As mobilization.

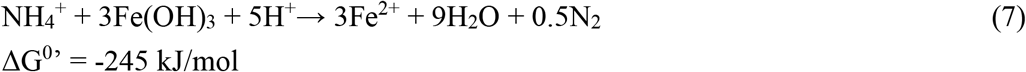

Moreover, Asammox-anaerobic ammonium oxidation coupled with As(V) reduction has been recently proposed in rice paddy soils (Zhang et al., 2022). This process could potentially increase the mobility of As since trivalent As is known to be generally more mobile than pentavalent As which tends to be easily adsorbed to Fe(III) minerals.

Ammonium, besides being produced by organic matter mineralization, can also originate from dissimilatory nitrate reduction to ammonium (DNRA) (equation 8). Many microorganisms from anoxic sediments can obtain energy via DNRA (Pandey et al., 2020). More importantly, N-DAMO archaea have also been shown to couple DNRA with CH_4_ oxidation (equation 9) suggesting that anaerobic CH_4_ oxidation might be coupled with NH_4_^+^ production (Nie et al., 2021). It is particularly relevant when the DOC/NO_3_^-^ molar ratio is high, then DNRA can replace denitrification as groundwater’s main NO_3_^-^ reduction pathway (Plummer et al., 2015).

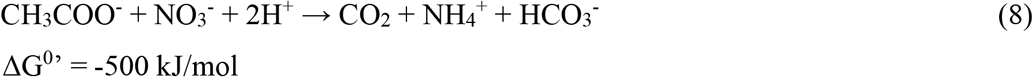

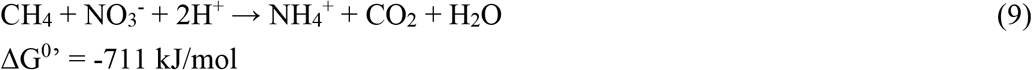

Additionally, when NO_3_^-^, NO_2_^-^ and NH_4_^+^ coexist in the redox interface, NO_3_^-^ and NH_4_^+^ can be converted into N_2_ together through the anammox process (equation 10).

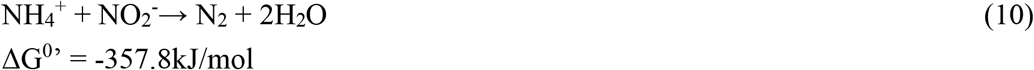

To date, all identified anammox bacteria belong to the order *‘Candidatus* Brocadiales’ within the phylum *Planctomycetes (Planctomycetota*) (Suarez et al., 2022). By conducting high-throughput sequencing of samples from aquifers around the world, Wang et al. estimated that anammox bacteria might be responsible for 80% of NO_3_^-^ and NO_2_^-^ removal at the global scale in these ecosystems (Wang et al., 2020).

Various N species can interact directly or indirectly with Fe(III) minerals. However, still very little is known about how the biological (trans)formation of N in an aquifer can affect the mobility of Fe and in consequence As. The input of N from the intensive application of fertilizers into methanogenic aquifers may stimulate N-DAMO processes, while DNRA may lead to the accumulation of NH_4_^+^ and potentially Feammox. Therefore, our present work aimed to assess the potential for anaerobic CH_4_ oxidation coupled to NO_3_^-^ reduction in As-contaminated aquifer sediments, evaluate the transformation pathways of NO_3_^-^, and investigate the possibility of Feammox potentially leading to As mobilization to groundwater. For this purpose, an As-bearing Fe(III)-rich sediment from an aquifer in Northeast Vietnam was anoxically incubated with ^13^CH_4_ and supplemented with ^15^NO_3_^-^. Additionally, the potential effect of N-DAMO on Fe and As (im)mobilization was studied by inoculating the sediment with N-DAMO enriched laboratory cultures. We monitored Fe speciation, CH_4_ and ^13^CO_2_ concentrations as well as N species evolution over time. Furthermore, to assess the composition of the microbial community we performed 16S rRNA gene amplicon sequencing at the end of the experiment.

## MATERIALS AND METHODS

### Study site and sediment sample collection

The study site is located in a rural area of the Red River delta, in Van Phuc village, about 15 km away from Hanoi, Vietnam (20°55’18.7’’N, 105°53’37.9?). The area’s geological, hydrochemical, and mineralogical characteristics have been studied previously (Berg et al., 2008; Eiche et al., 2017; Postma et al., 2017; Stopelli et al., 2020). In brief, the north-western part of the studied area is characterized as a Pleistocene aquifer consisting of brownish-orange sands and groundwater with low dissolved Fe^2+^ (less than 0.5 mg/L) and As concentrations <10 μg/L (Fig. 2). The south-eastern part consists of younger grey Holocene sands and As groundwater concentrations varying between 200-600 μg/L often surpassing the WHO standard of 10 μg/L by a factor of 10 to 50. The concentration of dissolved Fe^2+^ is also high (10 to 20 mg/L) indicating strongly reducing conditions (van Geen et al., 2013). Furthermore, the Holocene aquifer is characterized by nearly flammable CH_4_ concentrations (>50 mg/L) (Postma et al., 2017; Stopelli et al., 2021).

**Fig. 2.**
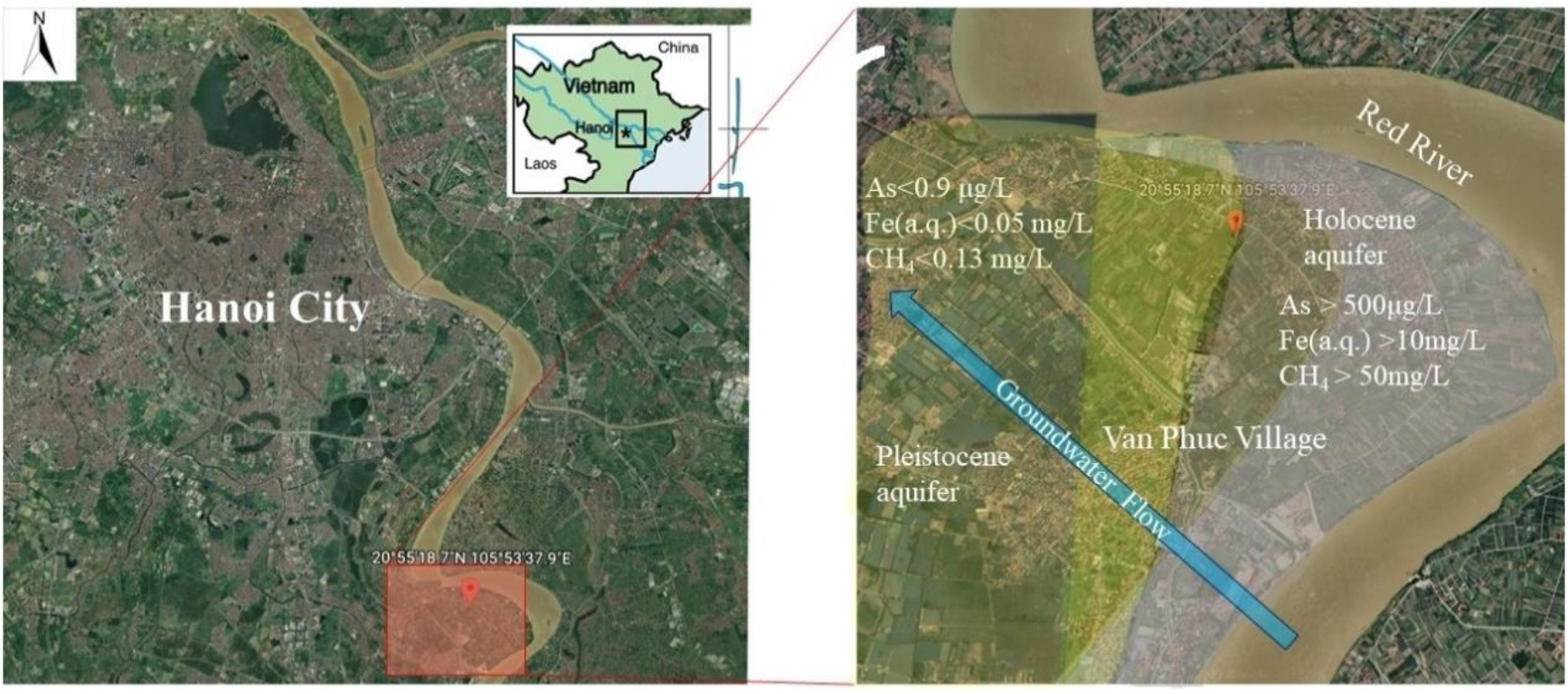
Satellite image of the study area. (red square in the left image). Zoom in to the drilling site located in the redox transition zone at the interface of the Pleistocene and Holocene aquifer (right). Google Earth, Maxar Technologies

In November 2018, a drilling campaign took place and sediment cores (9 cm diameter) were collected by rotary drilling up to 46.5 m below the ground, at the redox transition zone (RTZ) located in the interface of Holocene As-contaminated and Pleistocene pristine aquifer. The RTZ is subjected to intense geochemical and microbial activities which are suggested to be responsible for the As release to groundwater. Sediment samples were collected in water- and air-tight zip log bags (LamiZip, DAKLAPACK) with high barrier properties against oxygen and water vapor and protection against UV radiation to minimize sample alteration. All samples were flushed with N_2_ immediately after sampling and cooled during transportation to minimize microbial activity. Afterward, all samples were stored at 4°C anoxically in the dark until further use.

### Microcosms setup and incubation

For microcosm setups, the yellow-orange, less reduced sediment from 31m depth was used as this type of sediment is known to have a higher content of Fe(III) minerals and As compared to the grey reduced sediment. The sediment from this depth is characterized by Kontny et al. (2021). Microcosms were set up in 120 mL sterile glass serum bottles filled with 25 g of sediment and 50 mL synthetic groundwater medium (modified from Rathi et al. (2017); without As and Fe). Five different treatments were prepared in triplicates (Table 1): (1) amended with 0.2 ± 0.004 g (dry weight) biomass of N-DAMO(O) enrichment culture, 5 mM Na^15^NO_3_ (final concentration) and 0.8 mM ^13^CH_4_; (2) abiotic control – the same composition as treatment 1 with additional 150 mM of sodium azide (NaN_3_) to inhibit microbial activity; (3) amended with 0.2 ± 0.004 g (dry weight) of N-DAMO(V) enrichment culture, 5 mM Na^15^NO_3_ and 0.8 mM ^13^CH_4_; (4) only amended with 5 mM Na^15^NO_3_ and 0.8 mM ^13^CH_4_; (5) control group without any amendment. The N-DAMO(O) enrichment culture was obtained from an agricultural ditch in Ooijpolder, The Netherlands, and currently consists of *Ca*. Methanoperedens nitroreducens (~44%), *Ca*. Methylomirabilis (~26%) (Raghoebarsing et al., 2006; Schoelmerich et al., 2022). The N-DAMO(V) culture was enriched from rice paddy soil from Vercelli, Italy, and consists mainly of *Ca*. Methanoperedens (~78%) (Schoelmerich et al., 2022; Vaksmaa et al., 2017). Both cultures are grown in a continuous bioreactor under anoxic conditions with NO_3_^-^ as electron acceptor and CH_4_ as electron donor. Synthetic groundwater, Na^15^NO_3_, and NaN_3_ solution were gassed with N_2_/CO_2_ to remove dissolved O_2_ before use. All microcosms were prepared in an anoxic glovebox (97% N_2_ and 3% H_2_), closed with rubber stoppers and aluminum caps. The headspace gas was exchanged with N_2_/CO_2_ (9:1) until the final pressure of 1.83±0.05 bar to ensure CH_4_ dissolution and anoxic conditions. Afterward, microcosms were kept in the dark at 30°C and shaken at 30 r.p.m. for 64 days.

**Table 1.** Overview of microcosms setups used in the experiment

| Treatment | NAME | Inoculum | $^{15}\text{NO}_3$ | $^{13}\text{CH}_4$ | $\text{NaN}_3$ |
| --- | --- | --- | --- | --- | --- |
| 1 | N-DAMO(O) | N-DAMO(O) | ✓ | ✓ | x |
| 2 | Abiotic N-DAMO(O) | N-DAMO(O) | ✓ | ✓ | ✓ |
| 3 | N-DAMO(V) | N-DAMO(V) | ✓ | ✓ | x |
| 4 | $\text{NO}_3^- + \text{CH}_4$ | x | ✓ | ✓ | x |
| 5 | Control | x | x | x | x |

### Geochemical analyses

At each time point (day 0, 3, 6, 7, 15, 16, 24, 25, 33, 36, 41, 44, 52, 63), 2 mL of slurry were withdrawn by using a sterile syringe and needle (ø 1.20 × 40 mm) in an anoxic chamber. Samples were centrifuged at 14000 x g for 1 minute. Afterward, 100 μL of the supernatant was mixed with 100 μL 1M HCl to stabilize and dilute the sample for further Fe(II) quantification. One milliliter of supernatant was stabilized in 9 ml of 1% HNO_3_ for As, Fe, Mn, analysis by ICP-MS (8900, Agilent Technologies, USA). The remaining supernatant was transferred into an Eppendorf tube for NO_3_^-^, NO_2_^-^, and NH_4_^+^ quantification. Sediment was then digested with 1 mL 6M HCl for 1h, centrifuged for 5 min at 14000 r.p.m, and 100 μL of the supernatant was diluted with 100 μL 1M HCl. The concentrations of Fe(II) and total Fe were detected by the Ferrozine assay (Schaedler et al., 2018). The Griess assay was used to quantify NO_3_^-^ and NO_2_^-^ while the OPA assay was used to determine the concentration of NH_4_^+^ (Meseguer-Lloret et al., 2002; Sun et al., 2003).

The concentrations of ^13^CO_2_, ^15^N_2_O, and the ratio of ^30^N_2_/^28^N_2_ were determined by gas chromatography coupled to mass spectrometry (Trace DSQ II, Thermo Finnigan, Austin TX, USA), and the concentration of CH_4_ was quantified by gas chromatography with flame ionization detection (Hewlett Packard HP 5890 Series II Gas Chromatograph, Agilent Technologies, California, USA). Air pressure was also monitored at each sampling point by a portable pressure meter (GMH 3100, GHM Messtechnik, Regenstauf, Germany). The concentration of ^13^CO_2_ and ^15^N_2_O in the headspace and the total amount of ^13^CO_2_ and ^15^N_2_O in the incubation bottles were calculated following the formulas S1 and S2, respectively (Supplementary Materials).

### DNA extraction and microbial community analysis

The sediment sample for the DNA extraction was collected from the original sediment used for the incubation and from microcosms at the end of incubation (65 days). The DNA extraction was performed using the PowerSoil DNA extraction kit (DNeasy PowerSoil Pro Kit, QIAGEN, Hilden, Germany), according to the manufacturer’s protocol with an additional 500 μL of 10% (w/v) sterilized skim milk solution to the sediment sample, which may improve the DNA extraction yield from soil samples (Hoshino and Matsumoto, 2005). The concentration of the DNA was quantified using Qubit^®^ 2.0 Fluorometer with DNA HS kits (Life Technologies, Carlsbad, CA, USA). 16S rRNA gene amplicon sequencing was done by Macrogen (Macrogen, Amsterdam, The Netherlands) using the Illumina MiSeq Next Generation Sequencing platform. Paired-end libraries were constructed using the Illumina Herculase II Fusion DNA Polymerase Nextera XT Index Kit V2 (Illumina, Eindhoven, Netherlands). Primers used for bacterial amplification were Bac341F (5’-CCTACGGGNGGCWGCAG-3’; (Herlemann et al., 2011) and Bac806R (5’-GGACTACHVGGGTWTCTAAT-3’; (Caporaso et al., 2012). Archaeal amplification was performed with primers Arch349F (5’-GYGCASCAGKCGMGAAW-3’) and Arch806R (5’-GGACTACVSGGGTATCTAAT-3’; (Takai and Horikoshi, 2000). For Bacteria and Archaea separately, reads were trimmed and removed based on quality (settings: left trim 17 and 20, truncation length 267 and 270, maxE 2), followed by denoising and dereplication (settings: error learning bases 1e10, pooling during denoising, overhang trimming during merging) Amplicon Sequence Variant (ASV) calling, and finally taxonomic assignment (SILVA version nr138 training set, (Quast et al., 2013) and read abundance counting using DADA2 and its utilities (v1.22.0, (Callahan et al., 2016) in R (v4.1.2; R Core Team, 2019). After quality control and assignment of reads to ASVs, between 44679 and 115065 paired reads were assigned to 770 total archaea ASVs, and 44710 and 80477 paired reads were assigned to 944 total bacteria ASVs. Further analyses and visualization of these count and taxonomic data were performed also using R or Excel. The raw sequence data and metadata of the microcosms experiment have been deposited at the read sequence archive (SRA) database of the NCBI under the BioProject ID PRJNA887920.

## 3. RESULTS AND DISCUSSION

### Nitrogen species evolution

To assess the effect of NO_3_^-^ input on microbial processes in the Fe and As rich aquifers a microcosm experiment was conducted. Furthermore, we added N-DAMO cultures to the sediment incubations to monitor their effect on Fe and As mobilization. All microcosms supplemented with NO_3_^-^, except the abiotic one, efficiently removed nearly all NO_3_^-^ (5 mM) within 5 days of incubation. The abiotic control (treated with NaN_3_) showed no significant change in NO_3_^-^ concentration over time (Fig. 3a). It was somehow unexpected that in the uninoculated microcosms, NO_3_^-^ was reduced as efficiently as in N-DAMO inoculated microcosms. Previous studies from the same drilling site showed that the microbial community in sediment and water at 31 m depth has the potential for NO_3_^-^ reduction (Glodowska, 2021a; Glodowska et al., 2021). It was however surprising that the addition of N-DAMO cultures to the sedimentary native microbial communities showed a similarly high denitrification potential as native microbial community alone (equation 11). In the N-DAMO inoculated microcosms CH_4_, at least partially, served as an electron donor as decreasing NO_3_^-^ was positively correlated with decreasing ^13^CH_4_ concentration and increasing ^13^CO_2_ (Fig. 4a, b). However, in the inoculated as well as uninoculated microcosms, the native microbial community likely utilized recalcitrant organic C still present in the sediment for the heterotrophic NO_3_^-^ (Glodowska et al., 2020).

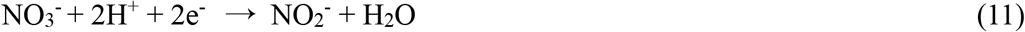

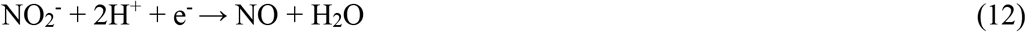

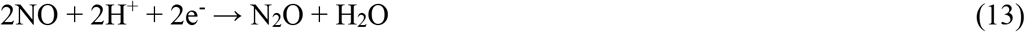

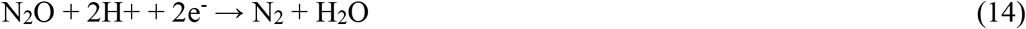

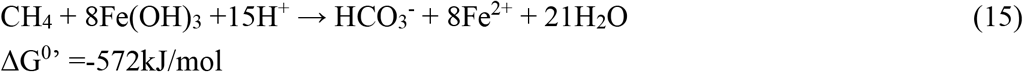

Although, the native microbial community in the setup supplemented with CH_4_ and NO_3_^-^ appeared to be capable of efficient NO_3_^-^ reduction, at a similar rate as the two N-DAMO inoculated setups, it, however, showed only a limited ability to further reduce NO_2_^-^ to other N-species (especially N_2_) (equation 12, 13, 14). In these microcosms NO_2_^-^ concentration rapidly increased to 1.12 mM within the first 2 days, dropped to 0.6 mM, and remained stable until the end of the experiment (Fig. 3b). Although our previous study showed that NC10 bacteria affiliating with *Ca*. Methylomirabilis that are known to reduce NO_2_^-^ at the expense of CH_4_ were present in the sediment and groundwater of this aquifer (Glodowska, 2021a, 2021b), their abundance in our experiment was probably too low to remove all NO_2_^-^. In the two N-DAMO inoculated treatments, NO_2_^-^ was nearly undetectable during the whole incubation time as both N-DAMO cultures can efficiently reduce NO_2_^-^. Interestingly, after supplying an additional 5 mM of NO_3_^-^ at the end of the experiment (after 65 days of incubation, Fig. S1) the native microbial community was dormant or appeared to have lost the ability to reduce NO_3_^-^. The tolerance of different microorganisms to NO_2_^-^ greatly varies (Guo and Gao, 2021), and the native denitrifying community may eventually have died due to prolonged exposure to relatively high concentrations of NO_2_^-^ (0.6 mM).

**Fig. 3.**
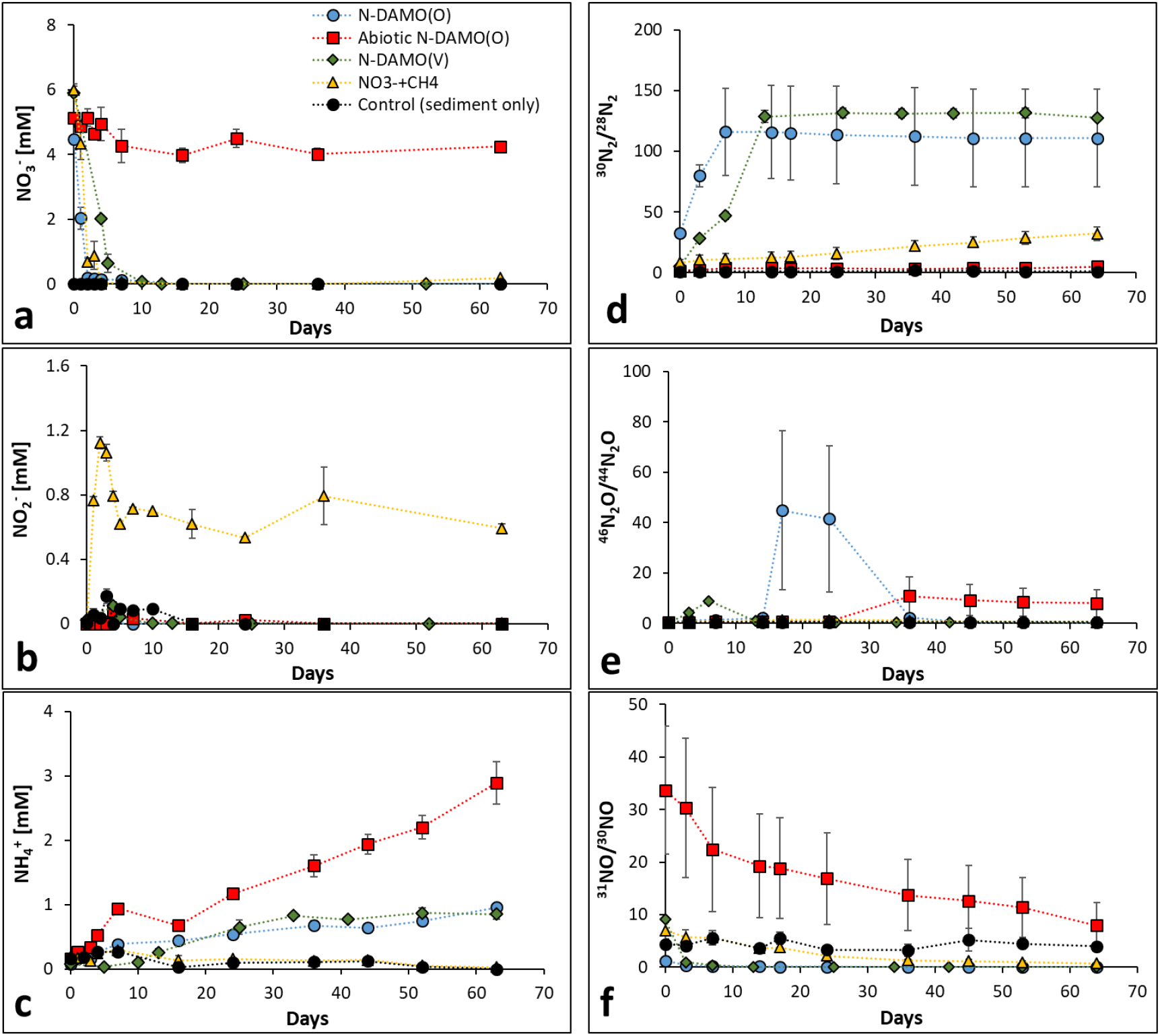
Evolution of N-compounds in microcosms within 65 days of aquifer sediment incubation. The concentration of a) NO_3_^-^, b) NO_2_^-^, and c) NH_4_^+^; and the ratio of d) ^30^N_2_/^28^N_2_, e) ^46^N_2_O/^44^N_2_O, and f) ^31^NO/^30^NO. Each microcosm was measured in technical triplicate, error bar stands for the standard deviation between biological triplicates of each treatment.

**Fig. 4.**
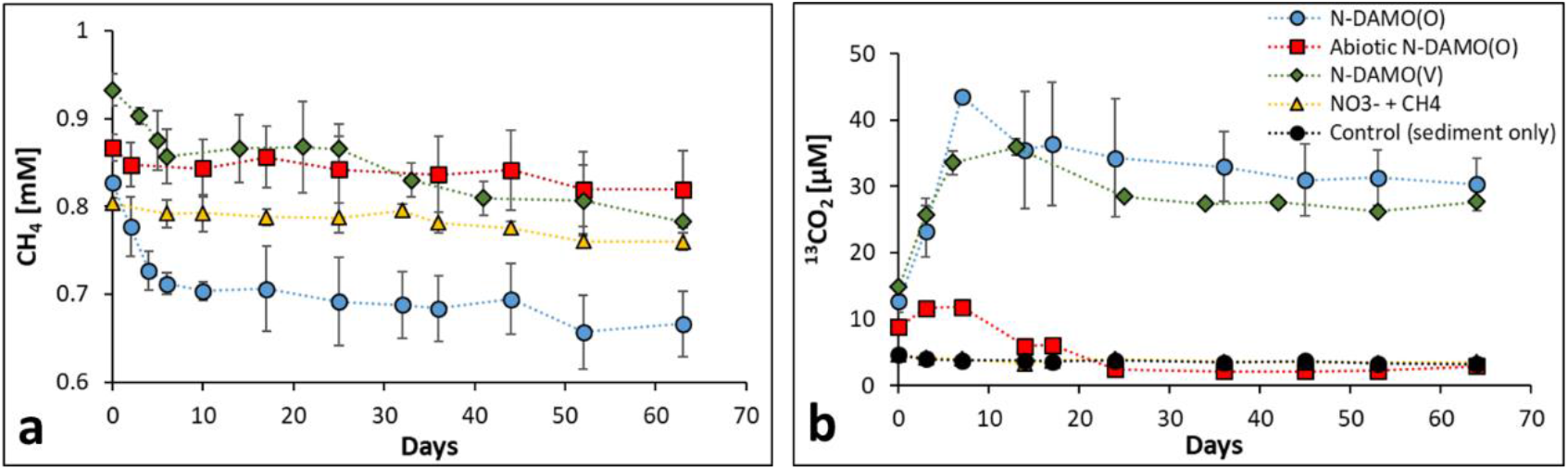
Methane oxidation and ^13^CO_2_ production in the five different microcosm setups. The concentration of (a) CH_4_ and (b) ^13^CO_2_ formation over time. For a better presentation of the changes in CH_4_ concentrations, the vertical axis starts at 0.6 mM as no CH_4_ was added to the control group. Only negligible methanogenesis was observed in the control setup (Fig. S5). Error bar stands for the standard deviation between biological triplicates of each treatment.

Denitrification is a stepwise process in which three intermediate species are produced; NO_2_^-^, NO, and N_2_O (equation 11, 12, 13) (Kuypers et al., 2018). In the NO_3_^-^ and CH_4_ supplemented microcosm, except for the already mentioned accumulation of NO_2_^-^, there was no significant accumulation of other N-intermediates and their concentration remained at very low levels till the end of the experiment (Fig. 3 e and f). Only the ratio of ^30^N_2_/^28^N_2_ in this group increased from 8 to 32% (Fig. 3 d). There was also no significant accumulation of NH_4_^+^ in the NO_3_^-^ and CH_4_ supplemented microcosm implying that neither mineralization of residual organic matter or dead biomass, nor considerable DNRA was taking place (Fig. 3c).

In the N-DAMO inoculated setups, N_2_ (Fig. 3d) and NH_4_^+^ (Fig. 3c) both began to increase immediately after the start of the experiment suggesting that both denitrification to N_2_ and DNRA were taking place. Specifically, in the N-DAMO(O) and N-DAMO(V) inoculated setups, the ^30^N_2_/^28^N_2_ ratio increased from 32 to 110% and from 5 to 127% at the end of the experiment, respectively (Fig. 3 d). In addition to ^30^N_2_, ^29^N_2_ was also generated in the N-DAMO(O) and N-DAMO(V) groups (reaching 2.4% and 2.2%, respectively) (Fig. S3), suggesting the activity of anammox bacteria. Two N-DAMO inoculated treatments converted about 20% of NO_3_^-^ to NH_4_^+^, which resulted in a NH_4_^+^ concentration of 0.95 and 0.85 mM, respectively, suggesting that DNRA is also an important way for N-DAMO cultures to reduce NO_3_^-^.

The concentration of NH_4_^+^ in the abiotic treatment increased considerably reaching almost 3 mM. This accumulation of NH_4_^+^ was most likely caused by the addition of NaN_3_, which is a strong bactericide, resulting in dying biomass and subsequent release of NH_4_^+^.

Although previous studies have pointed out that DNRA is an important source of NH_4_^+^ in aquifers (Weng et al., 2017), our results suggest that NH_4_^+^ does not originate from bacterial NO_3_^-^ reduction as NH_4_^+^ was generated only by the activity of inoculated N-DAMO enrichment cultures. Therefore, we suggest that the main source of NH_4_^+^ at the studied field site is the mineralization of buried organic matter and the potential infiltration of NH_4_^+^ from fertilizers. We also did not observe a parallel decrease in NH_4_^+^ concentration and an increase in Fe^2+^, therefore it is unlikely that Feammox takes place in the Van Phuc aquifer.

### CH_4_ oxidation and CO_2_ production

All the microcosms except a control setup were supplied with 10 mL ^13^CH_4_ (~ 0.8 mM). Only N-DAMO inoculated treatments however exhibited a considerable CH_4_ decrease over time. A particularly pronounced drop in CH_4_ concentrations was observed within the first 5 days of incubation which was also correlated with the formation of ^13^CO_2_ (Fig. 4a and b) and NO_3_^-^ reduction (Fig. 3a). It has to be beared in mind, however, that NO_3_^-^ reduction took place also in the uninoculated microcosms, therefore only part of the denitrification activity can be attributed to N-DAMO microorganisms. Specifically, the content of CH_4_ between day 0 and 6 decreased continuously in the N-DAMO(O) from 0.83 to 0.66 mM whereas in N-DAMO(V) CH_4_ dropped from 0.93 to 0.78 mM.

The CH_4_ oxidation was most pronounced in the N-DAMO(O) treatment which consumed 0.16 mM CH_4_ within 64 days (19.5 %). The increase of the ^45^CO_2_/^44^CO_2_ ratio in this group was also the highest, reaching nearly 18% after 7 days of incubation (Fig. S5). Although the amount of generated ^45^CO_2_ was similar between the two inoculated treatments (Fig. 4b), in the N-DAMO(V) inoculated microcosms the CH_4_ consumption was lower (0.15 mmol by day 64), and the ratio of ^45^CO_2_/^44^CO_2_ was also relatively low compared to the N-DAMO(O) treatment (Fig. S5). We calculated that the N-DAMO(O) treatment converted 21.9% of the consumed CH_4_ to CO_2_, while the N-DAMO(V) setup converted 13.7%. Despite the strong ability of N-DAMO enrichment cultures to couple CH_4_ oxidation to NO_3_^-^ reduction, less than 20% of NO_3_^-^ reduction in both N-DAMO groups was due to CH_4_ oxidation. The vast majority of NO_3_^-^ reduction is thus attributed to heterotrophic denitrification via oxidation of residual dissolved organic matter (DOM) still present within the sediment (Equations S3 and S4).

Although our previous study showed anaerobic CH_4_ oxidation coupled with Fe(III) reduction in this aquifer (Glodowska, 2020; Pienkowska et al., 2021), the CH_4_ concentration in uninoculated microcosms in this experiment remained stable until the end of incubation (65 days). This discrepancy is likely due to the much shorter incubation period of this experiment compared to our previous study, where Fe-DAMO activity was observed only after 100 days of incubation. It is also possible that Fe-DAMO was inhibited due to the presence of alternative electron acceptors such as NO_3_^-^. The Fe(III)-dependent CH_4_ oxidation could hovever take place after a longer period of incubation. Only a very small amount of methanogenic activity was observed within the orginal sediment as the CH_4_ concentration in the control incubations reached its maximum value of 2.13 μmol after 44 days, then dropped to ~1 μmol until the end of the incubation (Fig. S4).

### Iron reduction and As/Mn mobilization

All microcosms except the NO_3_^-^/CH_4_ supplemented uninoculated setups showed various degrees of Fe(III) reducing abilities (Fig 5). The most prominent Fe(III) reduction capacity was observed in N-DAMO inoculated setups, demonstrating that the N-DAMO enriched laboratory cultures have the potential to use Fe(III) as an electron acceptor. It has been previously shown that *Ca*. Methanoperedens species can indeed reduce Fe(III) (Cai et al., 2018; Ettwig et al., 2016) most likely due to the extraordinarily high number of *c*-type cytochromes (Kletzin et al., 2015; Leu et al., 2020). This implies that despite Fe(III) being a less favorable electron acceptor than NO_3_^-^ (equations 2 and 3), it can still be used to fuel the CH_4_ and/or DOM oxidation by N-DAMO, community members of the N-DAMO enrichment culture and/or the native microbial community. In our experiment, Fe(III) reduction was linked to organic carbon (OC) degradation, as we did not observe further CH_4_ oxidation and ^13^CO_2_ formation after NO_3_^-^ depletion. The native OM also stimulated Fe(III) reduction in uninoculated control microcosms as both dissolved and solid phase Fe(II) increased over time (Fig. 3a, b). The Fe(II) concentration in the abiotic setup and in the NO_3_^-^ and CH_4_ supplemented setup remained stable.

**Fig. 5.**
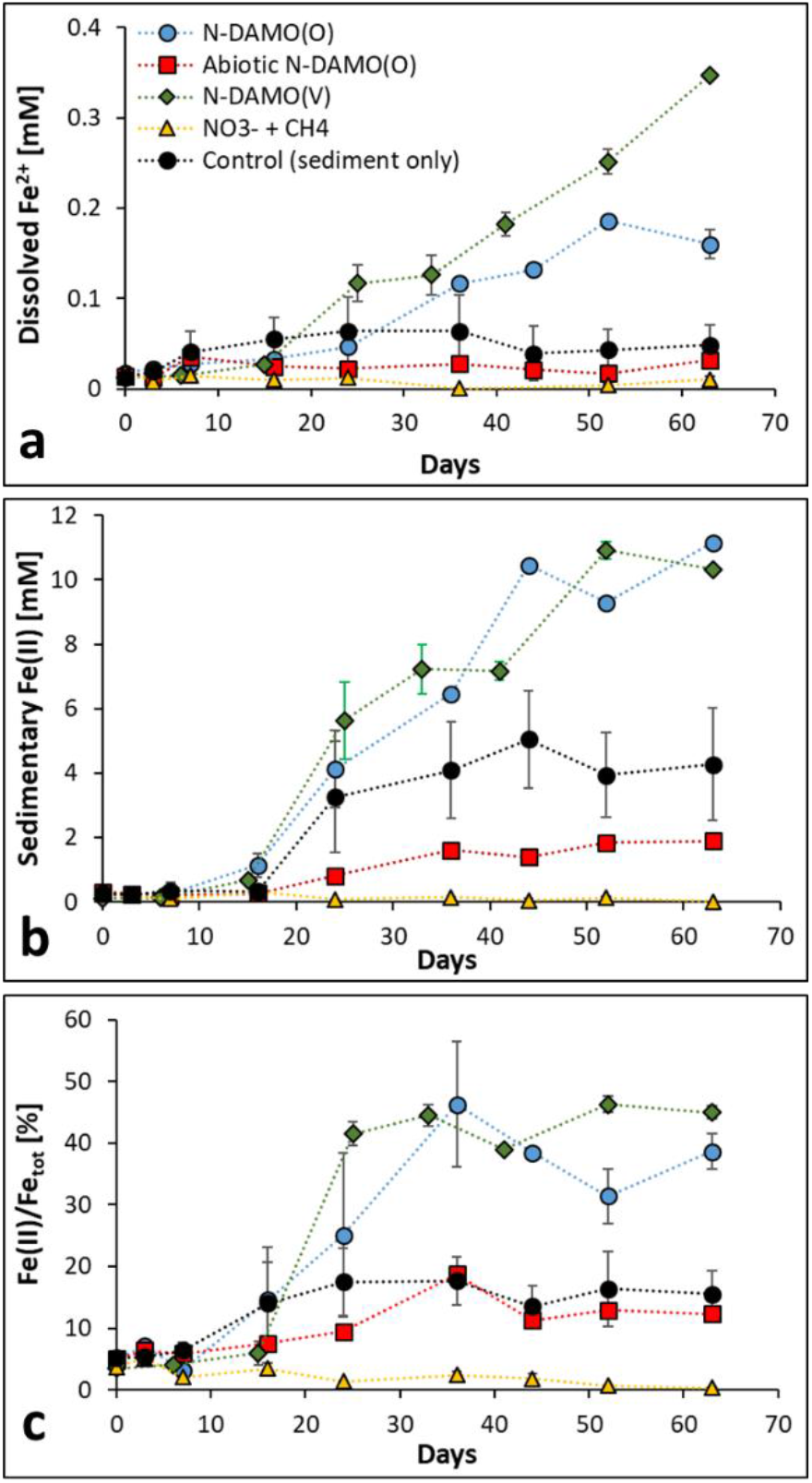
Changes in iron speciation. (a) Ferrous iron concentration in solution; (b) sedimentary Fe(II) concentration; and (c) Fe(II)/Fe_tot_ ratio. Error bar stands for the standard deviation between biological triplicates of each treatment.

Dissolved Fe^2+^ concentration in both N-DAMO setups started to rise after NO_3_^-^ depletion and the reduction rate increased significantly after 20 days, eventually reaching the highest values of 0.19 mM in N-DAMO(O) and 0.35 mM in N-DAMO(V) on days 52 and 64, respectively (Fig. 5a). However, a considerable amount of reduced Fe(III) remained as Fe(II) in the solid state (Fig. 5b). Fe(II) content in the sediment of the two N-DAMO inoculated treatments increased rapidly after depletion of NO_3_^-^, reaching a concentration of nearly 10 mM (Fig 5b). At the end of the experiment Fe(II) represented 39% and 45% of total Fe in N-DAMO(O) and (N-DAMO(V) inoculated setups, respectively (Fig. 5c).

The vast majority of Fe(II) in the two N-DAMO inoculated microcosms remained in the solid phase most likely because the aquifer’s sediment is rich in poorly crystalline Fe(III) minerals that are known to have a strong adsorption capacity for Fe(II) (Jeon et al., 2003). Furthermore, the formation of OM-Fe(II) complexes can also retain the newly formed Fe(II) (Du et al., 2018). In addition, CH_4_ oxidation produces a large amount of CO_2_, and a part of Fe^2+^ can return to the solid phase in the form of ferrous carbonate (FeCO3) (Appelo et al., 2002).

In the control microcosms, the native microbial community also showed a certain ability to heterotrophically reduce Fe(III) with native organic carbon serving as an electron donor. As there was no competition for electron acceptors due to the lack of NO_3_^-^ addition, the aqueous Fe^2+^ concentration in the control setup began to rise earlier than in the two N-DAMO inoculated setups (Fig 5a). Although some reduced Fe(II) was released as dissolved Fe^2+^ to the water phase, the majority of the Fe(II) remained sorbed in sediment (Fig. 5b). Overall, the ratio of Fe(II)/Fetot in control microcosms increased from 5 to 15.5% (Fig 5c).

Many previous studies have demonstrated that the addition of 150 mM NaN_3_ successfully inhibited the activity of the sedimentary microbial community (Cabrol et al., 2017; Glodowska et al., 2020), however, a minor increase in Fe(II) concentration in the here presented experiment indicated that it may have failed to completely inhibit N-DAMO’s Fe(III) reducing capability. According to previous studies, the bactericidal effectiveness of NaN_3_ is mainly due to inhibiting oxidative phosphorylation via inhibiting cytochrome oxidase (Harvey et al., 1999). However, it appears that under enhanced nutrient or anoxic conditions, the inhibitory effects of NaN_3_ might be reduced (Cabrol et al., 2017).

In NO_3_^-^/CH_4_ amended microcosms, no solid and aqueous Fe(II) formation was observed during the whole incubation period. This might be either because NO_3_^-^ was reduced as a more preferential electron acceptor exhausting bioavailable OC, and/or because of accumulation of NO_2_^-^ that potentially suppressed the metabolic activity of the native microbial community. Fang et al. investigated the effect of NO_3_^-^ on Fe(III) reduction and As release in the Datong Basin (Fang et al., 2021). Similarly, to our observations, they found that the presence of NO_3_^-^ can significantly inhibit the reduction of Fe(III) and thus decrease the release of As into the water. Unlike in the here presented experiment, Fang et al. used a pure culture of Fe(III) and NO_3_^-^ reducing *Bacillus* D2201, whereas we focused on microbial communities (native and N-DAMO enriched laboratory cultures). In our experiment, the reaction sequence followed the thermodynamical hierarchy of electron acceptors, i.e. Fe(III) reduction did not take place until NO_3_^-^ was depleted. However, in the Fang et al. experiments, the reduction of NO_3_^-^ and Fe(III) were carried out simultaneously. The inhibitory effects of NO_3_^-^ on dissimilatory Fe(III) reduction was also shown in a series of electron acceptor competition experiments with *Shewanella putrefaciens* (DiChristina, 1992). Overall, based on our experiments and previous observations, the presence of NO_3_^-^ clearly has an inhibitory effect on Fe(III) reduction.

The quantification of dissolved As suggested that Fe(III) reduction did not necessarily lead to As release (Fig. 6a). Only NO_3_^-^/CH_4_ amended microcosms did not show any Fe(III) reduction and As mobilization, whereas Fe(II) concentration in other treatments increased considerably. Fe(III) reduction was highest in the N-DAMO(V) inoculated treatment where surprisingly nearly no As was released from the sediment. Instead, a strong mobilization of Mn was observed. The highest dissolved Mn concentration here reached 33.5 mg/L after 25 days of incubation. Afterwards, the concentration of Mn started to decrease suggesting oversaturation and possible secondary mineral precipitation (Kawashima et al., 1988). The increasing dissolved Mn concentration was likely due to the reduction of Mn(IV) minerals, e.g. birnessite, present in the sediment by the N-DAMO(V) enrichment culture. It was previously shown that some *Ca*. Methanoperedens species (i.e *Ca*. Methanoperedens manganicus and *Ca*. Methanoperedens manganireducens) can reduce Mn(IV) to Mn(II) while oxidizing CH_4_ according to equation 15 (Leu et al., 2020).

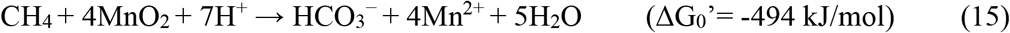

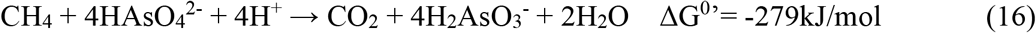

We suspect that *Ca*. Methanoperedens present in our N-DAMO(V) enrichment culture has a similar ability to use Mn(IV) as an electron acceptor.

**Fig. 6.**
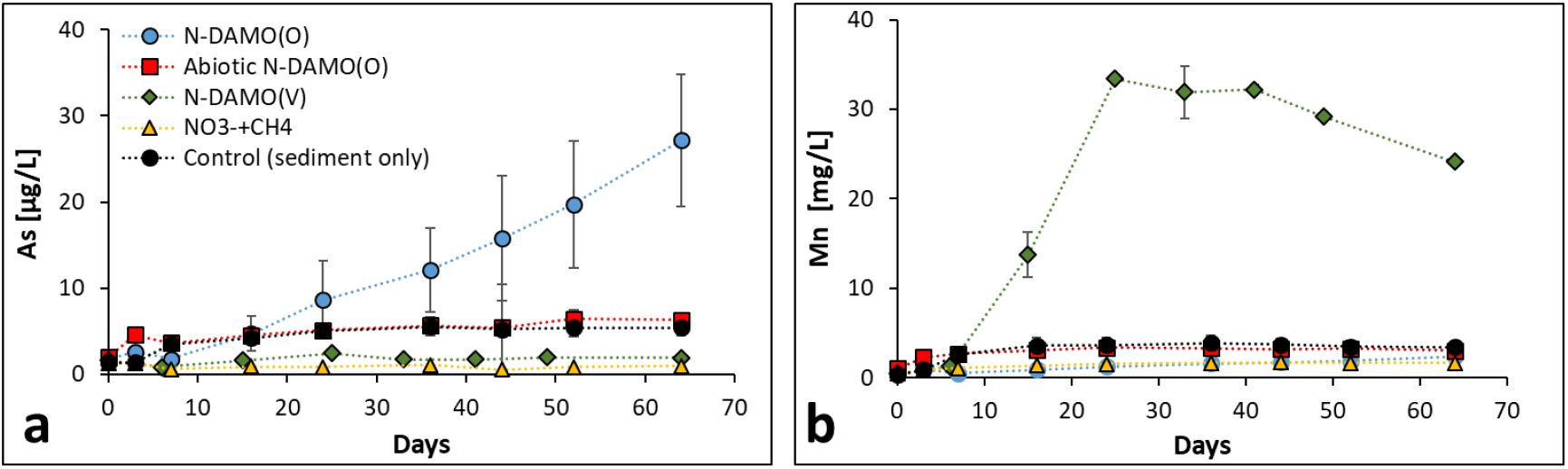
Change in concentration of (a) dissolved As and (b) Mn in water during 65 days of microcosm incubation. Please note that the concentration of Mn is in mg/L, and the concentration of As is in μg/L. Error bar stands for the standard deviation between biological triplicates of each group.

Inoculation with N-DAMO(O) led to high As mobilization reaching 27 μg/L at the end of the experiment, even though Fe(III) reduction was lower than in the treatment with added N-DAMO(V) enriched laboratory culture. The variation in As concentration between triplicates was, however, relatively large, suggesting that the dissolution of As is susceptible to environmental/abiotic factors. It has been suggested that some *Ca*. Methanoperedens archaea are genetically equipped to use As(V) as an electron acceptor as many of the available genomes encode for arsenate reductase (Arr) (Glodowska et al., 2022; Leu, 2020). However, until now there is a lack of laboratory studies linking the presence of *arr* genes with actual As(V) reduction and CH_4_ oxidation.

A study by Shi et al. demonstrated the coupling of anaerobic oxidation of CH_4_ with As(V) reduction in wetland soils (Shi et al., 2020). Metagenomic analysis in that study revealed, however, that the *arrA* gene was absent from ANME-2 metagenome-assembled genomes and instead found in non-methanotrophic *Sulfurospirillum* and *Geobacter*, therefore CH_4_ oxidation and the reduction of As(V) was likely mediated via a crossfeeding or syntrophic relationship of methanotrophic ANME archaea and As(V)-reducing bacteria. Nevertheless, it was concluded that CH_4_ oxidation coupled with As(V) reduction may contribute to up to 49% of As release in wetland soils. It is, therefore, possible that in our experiment As mobilization in N-DAMO(O) microcosms is not only due to reductive dissolution of As-bearing Fe(III) minerals but also via direct enzymatic reduction of As(V) by *Ca*. Methanoperedens, following equation 16 (Caldwell et al., 2008) or via metabolic collaboration between *Ca*. Methanoperedens and As(V) reducing bacteria.

### Changes in the microbial community

The archaeal 16S rRNA sequence abundance showed that the sediment used in the here presented experiment was dominated by *Ca*. Methanoperedens species (Fig. 7a). It is not surprising as our previous study also revealed high enrichment of this archaeon in the sediment (Glodowska et al., 2020). The native *Ca*. Methanoperedens however was likely very different from those present in enrichment cultures used here as it did neither show significant NO_3_^-^ reduction nor CH_4_ consumption in the uninoculated control setups. *Ca*. Methanoperedens was representing nearly 100% of the archeal community in the inoculated microcosms, while in NO_3_^-^/CH_4_ supplemented and control microcosms it accounted for 76 and 64%, respectively.

**Fig. 7.**
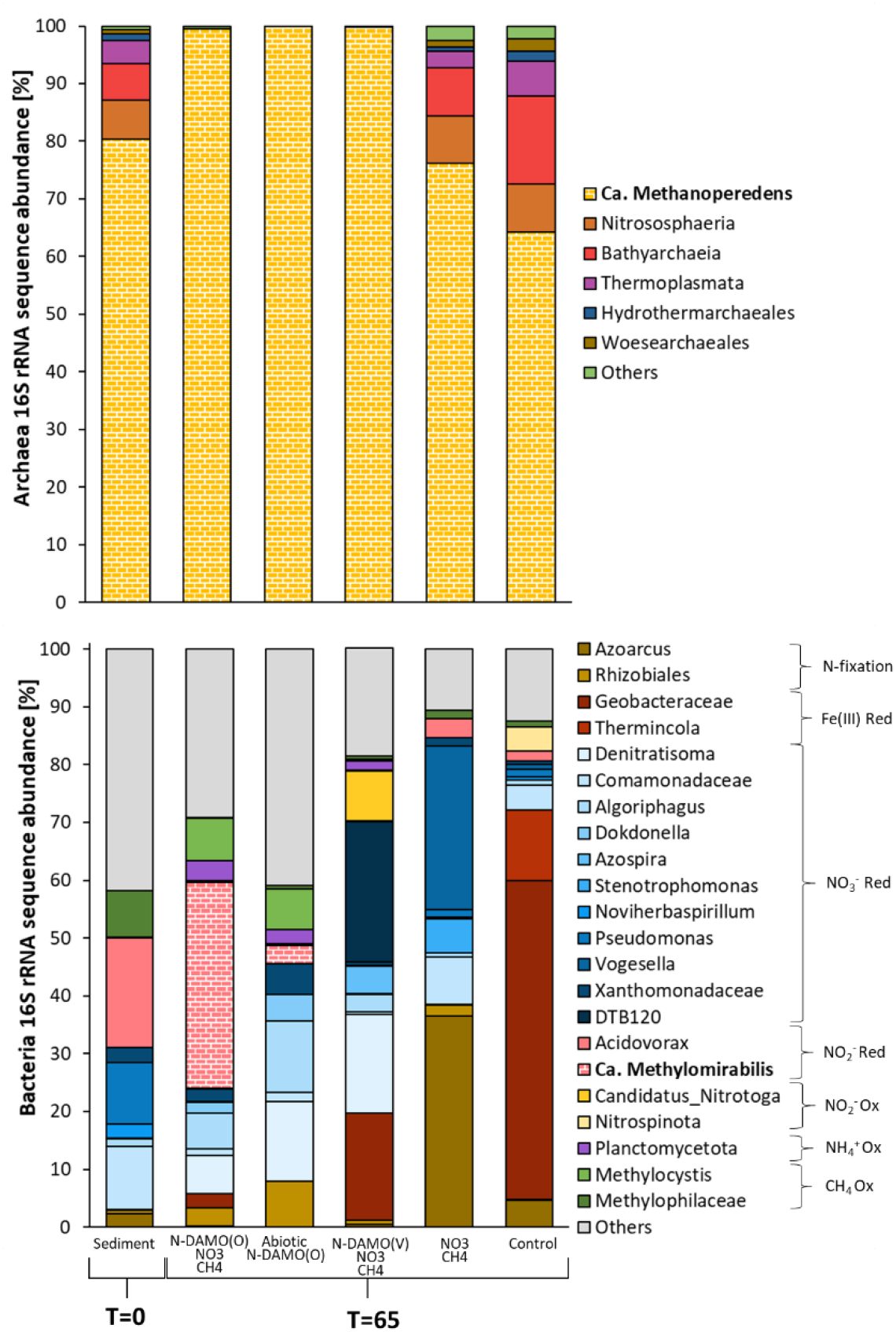
Relative abundance of archaeal and bacterial 16S rRNA gene sequence amplicons in microcosms at the end of the experiment (65 days). The highest taxonomic level is presented with its putative function. Sediment refers to the original sediment used in the experiment. Red = reduction, Ox = oxidation

The bacterial community substantially changed in composition over time (after 65 days of incubation) compared to the original sediment across all treatments. In the N-DAMO(O) inoculated microcosms *Ca*. Methylomirabilis was a dominant taxon (Fig. 7b). This is not surprising as *Ca*. Methylomirabilis represented nearly 30% of the N-DAMO(O) inoculum. In the abiotic N-DAMO(O) microcosms the relative abundance of *Ca*. Methylomirabilis was much lower, however, suggesting that introduced biomass was likely mineralized and their DNA degraded. In the N-DAMO(V) microcosms on the other hand bacteria from an uncharacterized phylum DTB120, named for an uncultured clone from a hot spring microbial mat (Lau et al., 2009), became the dominant taxon representing 24% of relative 16S rRNA gene abundance. It was previously suggested that these microorganisms might be involved in NO_3_^-^ reduction and Fe(II) oxidation (McAllister et al., 2021). Also, putative Fe(III)-reducers affiliating with Geobacteraceae increased in abundance, at the end representing over 18% of the bacterial community. Finally, *Denitratisoma*. a putative NO_3_^-^ reducer, also represented a substantial part of the microbial community reaching 17% relative abundance at the end of the experiment. The bacterial consortium in the NO_3_^-^/CH_4_ supplemented microcosms mainly consisted of denitrifying bacteria such as *Vogesella, Stenotrophomonas*, and bacteria within the *Comamonadaceae* family. Also, *Azoarcus*, a know N_2_-fixing bacterium was highly enriched at the end of the experiment (Zorraquino et al., 2018). Finally, the bacterial community in control microcosms was dominated by Fe(III)-reducing bacteria within the *Geobacteraceae* family and *Thermincola*, which is consistent with our previous observations (Glodowska et al., 2020).

## 4. Environmental Implications

Intensive use of N-fertilizers often comes with a risk of leaching and penetration of N compounds into aquifers thereby threatening groundwater quality. Moreover, the input of N compounds such as NO_3_^-^ which is a favorable electron acceptor will likely trigger microbiological processes and subsequently affect the hydrochemistry of water. Our study suggests that counterintuitively, the input of NO_3_^-^ to As-contaminated aquifer might be beneficial as NO_3_^-^ inhibits Fe(III) reduction and subsequently prevents As mobilization to groundwater. It suggests that NO_3_^-^ might be a more preferentially used electron acceptor than Fe(III) in Red River Delta sediment. This, however, might come with the risk of NO_2_^-^ production, as the native microbial community in the Van Phuc aquifer was not capable of rapid NO_2_^-^ reduction, leading to the accumulation of this toxic compound. Our study also demonstrated that although conditions appear suitable, the indigenous microbial community was not capable of N-DAMO yet input of NO_3_^-^ stimulated the denitrifying community. Although it was previously shown that Fe(III)-dependent anaerobic CH_4_ oxidation takes place in the Van Phuc aquifer, our study did not reveal parallel ^13^CO_2_ and Fe(II) formation. This is most likely due to the short incubation time (65 days) compared to the previous study (over 250 days). Finally, our geochemical data did not indicate potential anammox or Feammox activities. We did not observe a concomitant NH_4_^+^ decrease with the increase of Fe(II) concentration, it is however possible that the accumulation of NH_4_^+^ masked the consumption of NH_4_^+^ by Feammox.

The microcosm approach is a convenient way to screen the metabolic potential of native microbial communities, it can however not fully mimic the environmental conditions. For example, the leaching of fertilizers into groundwater is expected to be a slow and continuous supply of a rather low amount of NO_3_^-^. In our experiment, we supplied a single dose of 5 mM NO_3_^-^, which eventually led to the accumulation of NO_2_^-^ in the CH_4_/NO_3_^-^ amended treatment, inhibiting the activity of the native microbial community. To overcome this problem a dedicated bioreactor setup with a continuous supply of NO_3_^-^ could be designed. Further experiments are needed to fully understand the N cycling in groundwater aquifers and its impact on Fe and As mobility.

## Acknowledgments

This work was supported by the Netherlands Organisation for Scientific Research (N.W.O.) and Ministry of Education (OCW) through the Soehngen Institute of Anaerobic Microbiology Gravitation Grant 024.002.002. MJ was further supported by ERC Synergy MARIX 854088. The authors would like to thank all AdvectAs project members for their collaboration and support. Special thanks to Pham Hung Viet, Pham Thi Kim Trang, Vi Mai Lan, Mai Tran, and Viet Nga from Hanoi University of Science for their assistance during the sampling campaign.

