## Supplementary Materials for "Nitrate leaching and its implication for Fe and As mobility in a Southeast Asian aquifer"

### Calculation of the total amount of $^{13}CO_2$ and $^{15}N_2O$

$$\sum {}^{13}CO_2 = {}^{13}CO_{2(g)}\ [1 + kRT\ V_{liquid}/V_{gas}\ (1 + K_Z/[H^+])] \qquad \text{Equation S1}$$

Where:
1) $\sum{}^{13}CO_2$ is the total amount of $^{13}CO_2$ in the bottle
2) $^{13}CO_{2(g)}$ is the amount of $^{13}CO_2$ in the gas phase (headspace) in mmol.
3) k is the solubility coefficient of $CO_2$, which is $3.3x10^{-4}$ mol/m$^3$ Pa.
4) R is the universal gas constant, from the ideal gas law, which is 8.314 J mol$^{-1}$ K$^{-1}$.
5) T is the Kelvin temperature of the incubation condition. 4ºC = 277.15 K, RT is ~22 ºC = 295.15 K
6) $V_{liquid}$ and $V_{gas}$ are the volumes of the liquid and gas phases (in mL).
7) $K_Z$ is the dissociation constant of the first step of carbonic acid dissociation. That is $K_{a1}$, which you get from p$K_1$, which you have to calculate (see below).
8) $[H^+]$ is the molar concentration of $H^+$

The equation for calculating $N_2O$ is very similar to the equation S1

$$\sum {}^{14}N_2O = {}^{14}N_2O_{(g)}\ [1 + kRT\ V_{liquid}/V_{gas}] \qquad \text{Equation S2}$$

1) $\sum{}^{14} N_2O$ is the total amount of $^{14}$ $N_2O$ in the bottle
2) $^{14}$ $N_2O_{(g)}$ is the amount of $^{14}$ $N_2O$ in the gas phase (headspace) in mmol.
3) k is the solubility coefficient of $CO_2$, which is $2.4*10^{-4}$mol/m$^3$ Pa.
4) R is the universal gas constant, from the ideal gas law, which is 8.314 J mol$^{-1}$ K$^{-1}$.
5) T is the Kelvin temperature of the incubation condition. 4ºC = 277.15 K, RT is ~22 ºC = 295.15 K
6) $V_{liquid}$ and $V_{gas}$ are the volumes of the liquid and gas phases (in mL).
7) The $K_Z$ of $N_2O$ is 0 because it does not react with water.

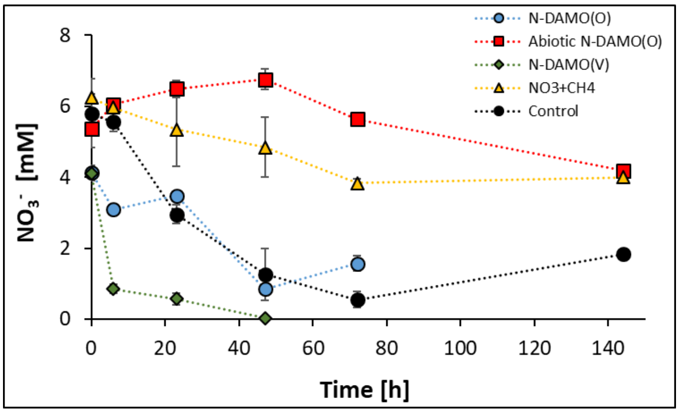


**Fig. S1 Changes of $NO_3^-$ concentration in five treatments at second $NO_3^-$ injection after 65 days of incubations**

Please note that the error bar in this figure represents the standard error of three measurements. One microcosm from each of the five treatments was used for this assay

After the second injection, the $NO_3^-$ concentration in the $CH_4$and $NO_3^-$ amended treatment only decreased by 2.24 mM in 150 hours (~ 6 days), which is much slower than the first $NO_3^-$ injection (almost completely depleted after 5 days). On the other hand, the native microbial community still possesses potent $NO_3^-$ reducing capability after 64 days of starvation.

This figure shows that the reducing ability of $NO_3^-$ and $NO_2^-$ in the native microbial community is greatly inhibited due to cytotoxicity caused by $NO_2^-$ accumulation.

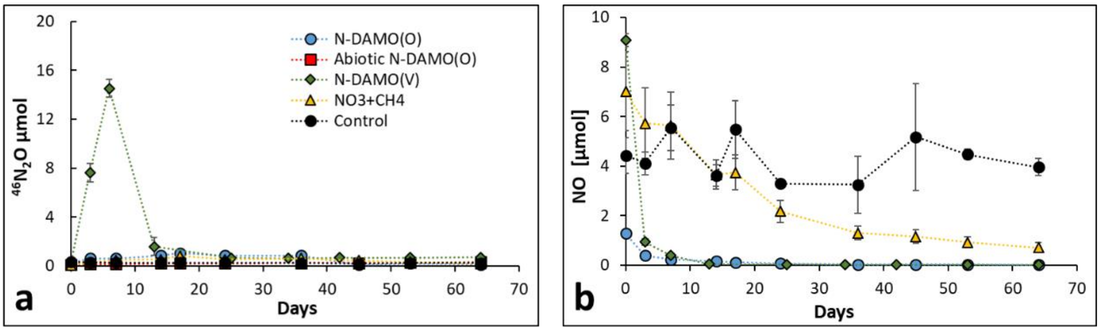


**Fig. S2 Changes in the contents of $^{46}N_2O$ a) and the content of NO b) ratio in the microcosms.**

According to Fig. S2a, only trace amounts of $^{46}N_2O$ were produced in the N-DAMO(V) group and were quickly consumed, while in other groups it remains at an extremely low level. As shown in Fig. S2b, the two N-DAMO enrichment cultures and native microbial communities exhibited the ability to consume NO, since the $^{31}NO/^{30}NO$ ratio was decreasing over time. The $^{31}NO/^{30}NO$ ratio is stable in the control group. Overall, we believe that the primary reduction products are $NH_4^+$ and $N_2$.

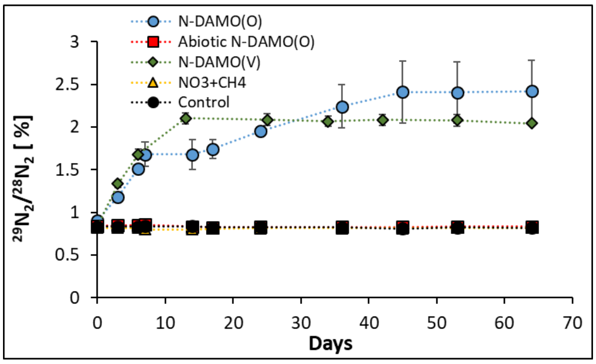


**Fig. S3 Changes in the ratio of $^{29}N_2$ to $^{28}N_2$ in the microcosms after 65 days of incubations.**

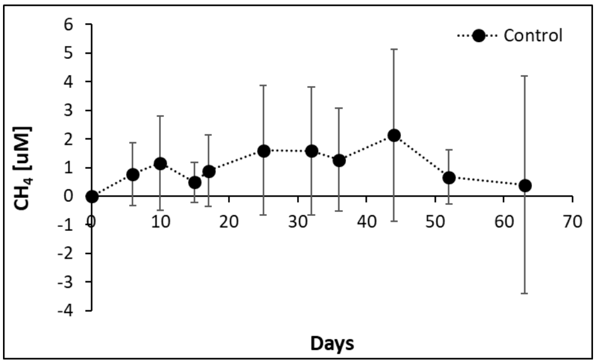


**Fig. S4 Changes in the concentration of $CH_4$ in the microcosms over the incubation time.** In addition to the information provided in Fig.2a, this figure shows that there is a weak methanogenic activity in the control group.

The concentration of $CH_4$ in the control group gradually increased from day 17 until it reached a maximum value of 2.13 μM on day 44 and then gradually decreased to 0.40 μM at the end of the incubation. In addition, the larger standard error also indicated that the intensity of methanogenesis activity varied greatly among the three replicas.

### Calculation of the contribution of $CH_4$ oxidation to $NO_3^-$ reduction.

$5CH_4 + 8NO_3^- + 8H^+ \rightarrow 5CO_2 + 4N_2 + 14H_2O$ Equation S3

According to formula S3, one molecule of $CH_4$ is oxidized, and the 1.6 $NO_3^-$ is reduced. Nitrate in the two N-DAMO inoculated groups was depleted in 10 days, and correspondingly, in N-DAMO(O) group, the $CH_4$ level dropped from 0.75 mmol to 0.72 mmol, and in N-DAMO(V) group, it decreased from 0.90 to 0.86 mmol, in the first 10 days of incubation.

We assume that all $CH_4$ consumption is due to the oxidation by $NO_3^-$. In the N-DAMO(O) group, where 30 μmol of $CH_4$ was oxidized, representing 48 μmol of $NO_3^-$

was reduced to $N_2$ or $NH_4^+$. Similarly, in N-DAMO(V) group, 40 μmol of $CH_4$ was consumed, meaning 64 μmol of $NO_3^-$ was reduced. At the beginning of the incubation, there were 310 μmol of $NO_3^-$ in the microcosms (5 mM*62 mL). From this, we can calculate that about 15.48% of total $NO_3^-$ reduction is attributed to $CH_4$ oxidation, whereas in N-DAMO(V) group this is 20.65%.

$$NO_3^- + CH_4 + 2H^+ \rightarrow CO_2 + NH_4^+ + H_2O \qquad \text{Equation S4}$$

In addition, according to formula S4, we also observed that N-DAMO enrichment culture undergoes the NDRA process, so for every molecule of $CH_4$ consumed, only one molecule of $NO_3^-$ is reduced to $NH_4^+$. In microcosms, the denitrification and DNRA processes were carried out simultaneously, and we do not know the contribution ratio of the two. We can speculate that the contribution of N-DAMO to the total $NO_3^-$ reduction in microcosms is lower than 20%, and the rest of the $NO_3^-$ reduction was caused by oxidation of DOM.

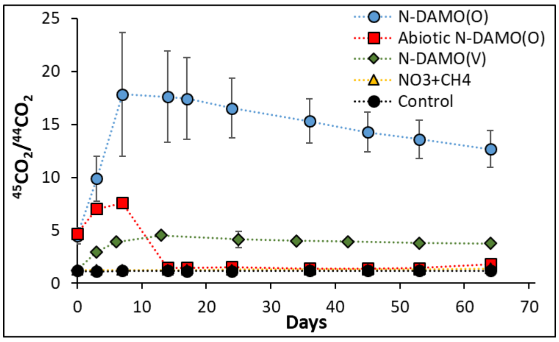


**Fig. S5 Changes in the content of $^{13}CO_2$ the microcosms,**
The content of $^{13}CO_2$ increased sharply in two inoculated groups, reaching 43.5 mmol on day 7 and 36 mmol on day 13 then gradually decreased, whereas the $^{13}CO_2$ content of the other three groups stay stable at around 3.5 mmol.
